# Accurate Prediction of Protein Complex Stoichiometry by Integrating AlphaFold3 and Template Information

**DOI:** 10.1101/2025.01.12.632663

**Authors:** Jian Liu, Pawan Neupane, Jianlin Cheng

## Abstract

Protein structure prediction methods require stoichiometry information (i.e., subunit counts) to predict the quaternary structure of protein complexes. However, this information is often unavailable, making stoichiometry prediction crucial for complexes with unknown stoichiometry. Despite its importance, few computational methods address this challenge. In this study, we present an approach that integrates AlphaFold3 structure predictions with homologous template data to predict stoichiometry. The method generates candidate stoichiometries, builds structural models for them using AlphaFold3, ranks them based on AlphaFold3 scores, and further refine predictions with template-based information when available. In the 16th community-wide Critical Assessment of Techniques for Protein Structure Prediction (CASP16), our method achieved 71.4% top-1 accuracy and 92.9% top-3 accuracy, outperforming other predictors in terms of the overall performance. This demonstrates the complementary strengths of AlphaFold3- and template-based predictions and highlights its applicability for uncharacterized protein complexes lacking stoichiometry data.

## 1. Introduction

Understanding the quaternary structure of protein complexes (multimers) is crucial for studying their function and has applications in protein design, engineering, and drug discovery. Computational tools like AlphaFold-Multimer^1,2^ and AlphaFold3^3^ have significantly advanced protein complex structure prediction but rely on prior stoichiometry information, i.e., the number of copies (count) of each unique subunit in a complex, to predict complex structures. This limitation excludes many complexes with unknown stoichiometry, underscoring the need to predict stoichiometry directly from sequences. Despite its importance, this problem remains underexplored, with few available methods^4,5^.

To address this gap, we propose a method that leverages AlphaFold3 to generate structural models for a range of stoichiometry candidates of a target complex, followed by ranking these candidates based on quality scores of the models. Given the computational cost of generating structural models, a manageable candidate list is essential. This can be achieved by utilizing known stoichiometry data from homologous templates in the Protein Data Bank (PDB) to define a feasible range of candidates. For homo-multimers (complexes with multiple identical subunits), generating structural models by enumerating subunit counts within AlphaFold3’s capacity often suffices, while template information can further narrow the options.

This method also assumes AlphaFold3 ranking scores for correct stoichiometries are higher for incorrect ones, which is not always guaranteed. To improve accuracy, confident template-based stoichiometry predictions are used to refine rankings when available. Combining these elements, we developed **PreStoi**, a novel approach integrating AlphaFold3 predictions, homologous template data, and template-based stoichiometry adjustment.

PreStoi was employed by our MULTICOM predictors in the 2024 community-wide CASP16 experiment, where participants predicted protein complex structures without prior stoichiometry information. Accurate stoichiometry prediction was critical, as errors significantly reduced structural prediction accuracy.

We compared MULTICOM predictors’ stoichiometry predictions to those of other CASP16 predictors whose stoichiometry prediction methods were described in the CASP16 abstract book. The results showed that PreStoi outperformed all other predictors, demonstrating its ability to accurately predict stoichiometry and marking a significant advancement in the field. As a robust tool for analyzing uncharacterized protein complexes, PreStoi also lays the groundwork for developing more advanced stoichiometry prediction methods.

### 2. Materials and Methods

The workflow of PreStoi, illustrated in **Figure 1**, consists of three steps: (1) template-based stoichiometry candidate generation and prediction, (2) AlphaFold3-based stoichiometry ranking and prediction, and (3) integration of AlphaFold3- and template-based stoichiometry predictions, detailed as follows:

**Figure 1.**
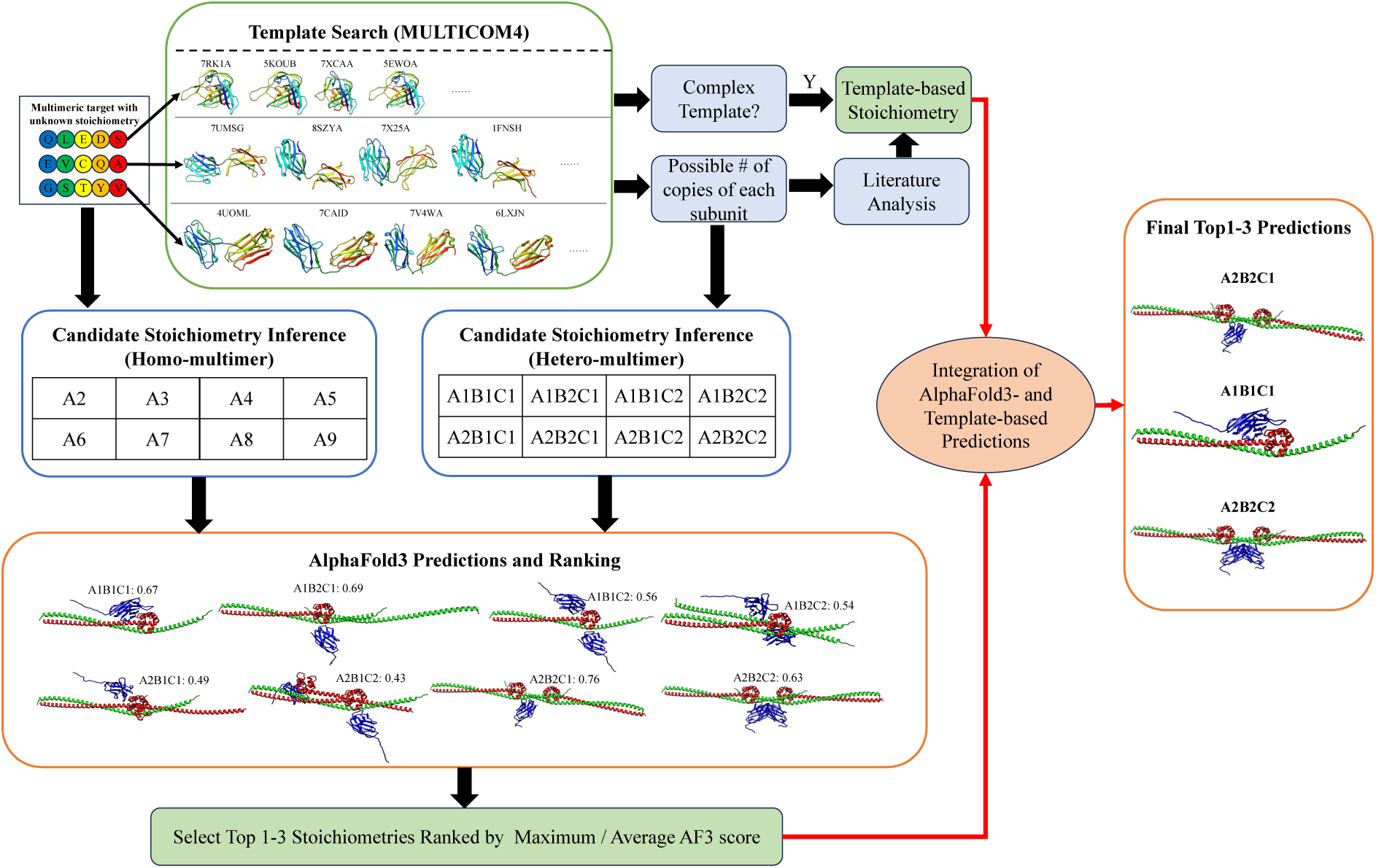
The workflow of predicting stoichiometry of protein complexes (multimers).

### 2.1. Template-based Stiochiometry Candidate Generation and Prediction

PreStoi begins by utilizing the enhanced MULTICOM4 pipeline (an upgraded version of MULTICOM3^6,7^) to identify structural templates for each unique subunit of a protein complex. The pipeline employs Jackhmmer^8^ to search each subunit sequence against Uniref90^9^, generating a multiple sequence alignment (MSA). This MSA is used by HHSearch^10^ to build sequence profiles, which are compared against the profiles of the proteins with known structures in Protein Data Bank^11^ (PDB) to find homologous templates.

Templates with known stoichiometry provide ranges for the number of copies of each subunit. These ranges are used to generate combinations of subunit copies, resulting in a list of stoichiometry candidates. For example, if a complex has three unique subunits (A, B, and C) and each subunit appears in one or two copies across templates, eight possible stoichiometries (i.e., A1B1C1, A1B1C2, A1B2C1, A1B2C2, A2B1C1, A2B1C2, A2B2C1, A2B2C1) (**Figure 1**) are generated. This approach significantly reduces the number of candidates needing evaluation.

For homo-multimers (identical subunits), the number of candidates is smaller than for hetero-multimers consisting of multiple different subunits. Simply enumerating subunit copies from two (dimer) to the maximum number AlphaFold3 can process—typically 2 to 9 in our experiments— suffices. Template information can further narrow candidates for homo-multimers.

In rare cases where no significant template is identified, the threshold for recognizing homologous templates can be relaxed to include less significant ones. Thus, a list of stoichiometry candidates can always be generated for further evaluation.

If all subunits in a complex share a common, significant complex template from the PDB (i.e., the same PDB code), the stoichiometry of that template is directly used as the prediction, referred to as *template-based stoichiometry prediction*. When no complex template covers all subunits, analyzing individual subunit templates, partial complex templates, and related literature may yield stoichiometry prediction. However, such analysis requires manual analysis and is not fully automated. Overall, the template-based stoichiometry prediction was applied to about 61% of CASP16 targets because the remaining targets did not have significant complex templates for making such prediction.

### 2.2. AlphaFold3-based Stoichiometry Ranking and Prediction

For each stoichiometry candidate, AlphaFold3 generates a set of structural models, typically 25–50 per candidate, with the exact number adjustable based on available computing resources. During the CASP16 experiment, models were generated using the AlphaFold3 webserver, as the source code was not yet released. This process has since been automated using the local version of AlphaFold3.

AlphaFold3 assigns a ranking score to each model, calculated as a weighted sum of several metrics, including predicted global structural quality (e.g., predicted TM-score^12^ or pTM), predicted interface quality (e.g., interface pTM or ipTM), a reward for disordered regions, and a penalty for structural clashes. The maximum or average ranking score of the models for each stoichiometry candidate (denoted as AF-max or AF-avg) is used to score the stoichiometry.

Candidates are ranked by their scores, and usually only the top 1–3 stoichiometries are selected for further analysis, as they almost always include the correct stoichiometry if it is present in the candidate list.

### 2.3. Integration of AlphaFold3- and Template-based Stoichiometry Predictions

If no template-based stoichiometry prediction is available, the selected AlphaFold3-based predictions are used as the final prediction. However, when template-based predictions are available, they are cross-checked with AlphaFold3-based predictions to enhance accuracy as follows.

If the top-1 AlphaFold3-predicted stoichiometry matches the template-based prediction, it is retained as the top choice. In cases of conflict, a confident template-based prediction—strongly supported by complex templates or literature evidence—takes precedence and is assigned as the top-1 prediction. The highest-ranked AlphaFold3 prediction is still included within the top three. Occasionally, weak template-based evidence or some features of AlphaFold3 structural models such as symmetry, stability, and interaction interfaces can be used to refine the AlphaFold3-based ranking of stoichiometries.

Typically, one to three final stoichiometry predictions are made for each target, depending on the confidence level, as indicated by the significance of templates or AlphaFold3 ranking scores. Occasionally, one or two more predictions can be added if the prediction confidence is very low.

### 2.4. CASP16 Multimer Targets for Stoichiometry Prediction

During Phase 0 of the CASP16 experiment, 28 complex targets (9 homo-multimers and 19 hetero-multimers) were released without stoichiometry information for protein structure predictors to predict their quaternary structures. Predictors were required to first predict the stoichiometry of each target before generating structural models based on the predicted stoichiometry.

Each CASP16 predictor could submit up to five structural models per target, which could include up to five different stoichiometries. The true stoichiometries of the 28 targets, listed in **Table 1**, were used to evaluate the performance of the MULTICOM predictors based on PreStoi, as well as other CASP16 predictors whose stoichiometry prediction methods were described in the CASP16 abstract book^13^. A stoichiometry prediction was considered correct only if it matched the true stoichiometry.

**Table 1.** The 28 CASP16 Phase 0 targets, comprising 9 homo-multimers and 19 hetero-multimers, along with their stoichiometry information. Unique subunits in each target are represented by letters (A, B, C, etc.), with the number following each letter indicating the number of copies for each subunit. For example, “A1B2” signifies a complex with two subunits: one copy of subunit A and two copies of subunit B.

| Homo-multimers |  | Hetero-multimers |  |
| --- | --- | --- | --- |
| Target # | Stoichiometry | Target # | Stoichiometry |
| T0206o | A2 | H0208 | A1B1 |
| T0218o | A2 | H0215 | A1B1 |
| T0234o | A3 | H0217 | A2B2C2D2E2F2 |
| T0235o | A6 | H0220 | A1B4 |
| T0237o | A4 | H0222 | A1B1C1 |
| T0240o | A3 | H0223 | A1B1C1 |
| T0257o | A3 | H0225 | A1B1C1 |
| T0259o | A3 | H0227 | A1B6 |
| T0270o | A6 | H0229 | A1B1 |
|  |  | H0230 | A1B1 |
|  |  | H0232 | A2B2 |
|  |  | H0233 | A2B2C2 |
|  |  | H0236 | A3B6 |
|  |  | H0244 | A2B2C2 |
|  |  | H0245 | A1B1 |
|  |  | H0258 | A1B2 |
|  |  | H0265 | A9B18 |
|  |  | H0267 | A2B2 |
|  |  | H0272 | A1B1C1D1E1F1G1H1I1 |

## 3. Results

### 3.1 Comparison of MULTICOM_AI and other CASP16 predictors in stoichiometry prediction

During CASP16, we blindly tested five predictors (MULTICOM, MULTICOM_AI, MULTICOM_GATE, MULTICOM_LLM, and MULTICOM_human), all of which utilized identical or slightly modified versions of PreStoi. These predictors achieved comparable performance. To streamline the comparison, we focus solely on the performance of one of our server predictors - MULTICOM_AI - relative to other CASP16 predictors. **Figure 2** compares the stoichiometry prediction performance of our MULTICOM_AI predictor, using PreStoi, against other 20 CASP16 predictors whose methods were detailed in the CASP16 abstract book (NKRNA-s^14^, Zheng-Multimer^15^, Schneidman^16^, Zheng^17^, MIEnsembles-Server^18^, CSSB_experimental^19^, CSSB-Human^20^, GromihaLab^21^, KiharaLab^22^, OpenComplex^23^, OpenComplex_Server^23^, MultiFOLD2^24^, McGuffin^25^, elofsson^26^, PEZYFoldings^27^, AF3-server^26^, kiharalab_server^22^, APOLLO^28^, ARC^29^, COAST^28^.).

**Figure 2.**
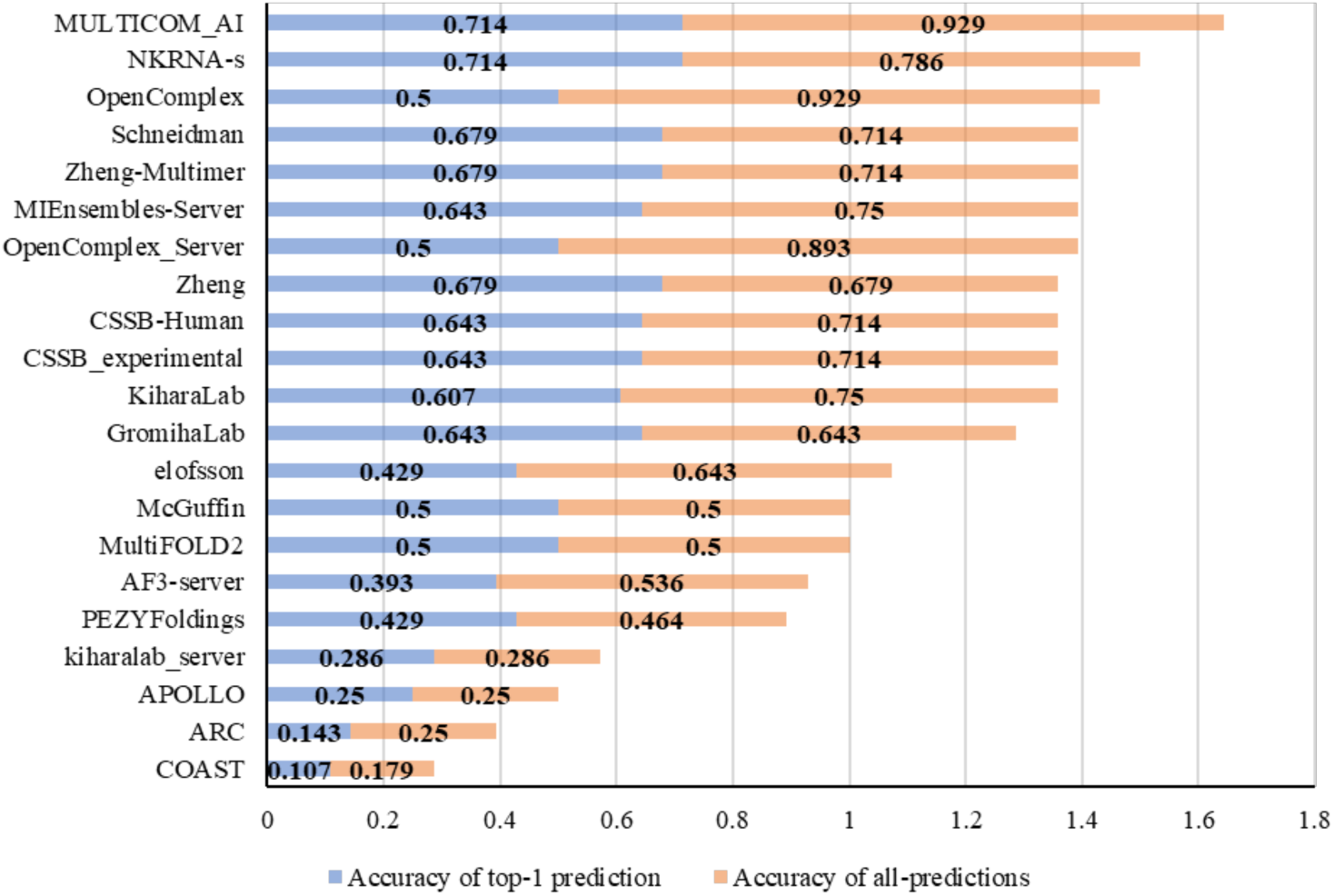
The stoichiometry prediction accuracy of the 21 CASP16 predictors on 28 Phase 0 targets.

Each CASP16 predictor submitted up to five structural models per target, with stoichiometry information extracted from the submitted models available on the CASP16 website. Although most predictors submitted five models per target, the number of unique stoichiometries per target was typically fewer than five. Prediction accuracy was assessed in two ways: *top-1 prediction accuracy*, which measures the percentage of targets where the top-ranked stoichiometry was correct, and *all-predictions accuracy*, which evaluates the percentage of targets where any submitted stoichiometry was correct.

As shown in **Figure 2**, MULTICOM_AI achieved the highest top-1 prediction accuracy of **71.4%** on the 28 CASP16 targets, matching NKRNA-s and surpassing all other predictors, whose accuracies ranged from 10.7% to 67.9%. For all-predictions accuracy, MULTICOM_AI achieved **92.9%**, substantially higher than NKRNA-s (78.6%) and tying with OpenComplex as the most accurate. However, OpenComplex had a much lower top-1 prediction accuracy (50%), indicating difficulty in ranking the correct stoichiometry as the top-1 prediction. Considering both top-1 and all-predictions accuracy, MULTICOM_AI clearly outperformed all other predictors (**Figure 2)**. The average of top-1 prediction accuracy and all-prediction accuracy of all the predictors is reported in supplemental **Table S1**).

To our knowledge, CASP16 was the first community benchmark for stoichiometry prediction. Consequently, most participants, including us, developed their methods during the competition. This likely contributed to the relatively low accuracy (<50%) of many predictors, which adversely affected their Phase 0 complex structure predictions. Incorrect stoichiometry predictions often led to significantly lower-quality structural models than correct ones.

The superior stoichiometry prediction accuracy of PreStoi contributed to our MULTICOM predictors’ success of ranking no. 1 in Phase 0 complex structure prediction at CASP16 (see CASP16 Phase 0 results: https://predictioncenter.org/casp16/results.cgi?tr_type=multimer&phase=0&groups_id=&model=1), outperforming all other CASP16 complex structure predictors. This demonstrates the value of the accurate stoichiometry prediction method developed in this work, which can benefit other protein structure prediction efforts within the community.

### 3.2 Analysis of the overall performance of MULTICOM_AI in the CASP16 experiment

**Table 2** summarizes the stoichiometry prediction results of PreStoi, implemented within our MULTICOM_AI protein structure predictor, on 28 CASP16 multimer targets. All stoichiometries predicted by MULTICOM_AI during CASP16 are detailed in supplemental **Table S2**. MULTICOM_AI predicted up to three stoichiometries per target, typically only one or two, except for three targets T0206o, H0208 and H0244, where more than three stoichiometries were predicted. For consistency, only the top three stoichiometry predictions for them were included in the evaluation.

**Table 2.** Accuracy and results of MULTICOM_AI stoichiometry prediction on 28 CASP16 targets. Target difficulty is measured as the percentage of CASP16 predictors that made incorrect top-1 predictions for each target, with higher percentages indicating greater difficulty. Targets with a difficulty value below 60% are classified as easy targets (green background), while those with 60% or higher are classified as hard targets (orange background). AF-max and AF-avg refer to the use of AlphaFold3’s maximum and average ranking scores, respectively, to determine the top-1 stoichiometry prediction. TB represents the template-based prediction. The Final Prediction (top 1) column reflects the combined top-1 prediction of MULTICOM_AI based on both AlphaFold3 and template-based approaches. The Decision Choice column describes which method was ultimately selected for the final top-1 prediction. The Top 3 column indicates whether any of the top 3 final predictions of MULTICOM_AI for a target was correct. The last two rows summarize the accuracy of each approach: AF-max (top 1), AF-avg (top 1), TB (top 1), Final Prediction (top 1), and Final Prediction (top 3), evaluated across all 28 targets or only those to which the respective method was applied, where the number of correctly predicted targets / the total number of targets considered in evaluating an approach is listed beneath its accuracy number. N/A signifies that a method was not applied to a target, either due to the target’s size (e.g., ≥5000 residues, which exceeded the AlphaFold3 web server’s capacity) or the absence of template-based predictions. Correct predictions are shown in red font. T####o denotes homo-multimer targets, while H#### represents hetero-multimer targets. Among the 28 targets, 9 are homo-multimers, and 19 are hetero-multimers.

| Target (28) | Difficulty | True stoichiometry | AF-max | AF-avg | TB | Final prediction (Top 1) | Decision choice | Final Top 3 |
| --- | --- | --- | --- | --- | --- | --- | --- | --- |
| H0225 | 10% | A1B1C1 | A1B1C1 | A1B1C1 | A1B1C1 | A1B1C1 | AF-max + AF-avg + TB | Yes |
| H0222 | 12% | A1B1C1 | A1B1C1 | A1B1C1 | A1B1C1 | A1B1C1 | AF-max + AF-avg + TB | Yes |
| H0223 | 13% | A1B1C1 | A1B1C1 | A1B1C1 | A1B1C1 | A1B1C1 | AF-max + AF-avg + TB | Yes |
| H0245 | 15% | A1B1 | A1B1 | A1B1 | N/A | A1B1 | AF-max + AF-avg | Yes |
| H0215 | 25% | A1B1 | A1B1 | A2B2 | A1B1 | A1B1 | AF-max + TB | Yes |
| T0206o | 28% | A2 | A2 | A2 | A2 | A2 | AF-max + AF-avg + TB | Yes |
| T0257o | 28% | A3 | A3 | A2 | A3 | A3 | AF-max + TB | Yes |
| T0259o | 29% | A3 | A4 | A3 | N/A | A3 | AF-avg | Yes |
| H0208 | 34% | A1B1 | A1B1 | A1B1 | A2B2 | A2B2 | TB | Yes |
| T0240o | 37% | A3 | A3 | A2 | N/A | A3 | AF-max | Yes |
| H0232 | 43% | A2B2 | A2B2 | A2B2 | A2B2 | A2B2 | AF-max + AF-avg + TB | Yes |
| H0272 | 43% | A1B1C1D1E1<br>F1G1H1I1 | N/A | N/A | A1B1C1D1E1<br>F1G1H1I1 | A1B1C1D1E1<br>F1G1H1I1 | TB | Yes |
| H0220 | 47% | A1B4 | A1B1 | A1B1 | A1B4 | A1B4 | TB | Yes |
| T0237o | 59% | A4 | A4 | A4 | N/A | A4 | AF-max + AF-avg | Yes |
| H0233 | 62% | A2B2C2 | A1B1C1 | A1B1C1 | A2B2C2 | A2B2C2 | TB | Yes |
| H0227 | 63% | A1B6 | N/A | N/A | A1B6 | A1B6 | TB | Yes |
| T0235o | 63% | A6 | A6 | A6 | N/A | A6 | AF-max + AF-avg | Yes |
| T0234o | 66% | A3 | A3 | A3 | N/A | A3 | AF-max + AF-avg | Yes |
| H0229 | 69% | A1B1 | A1B1 | A2B2 | N/A | A2B2 | AF-avg | Yes |
| H0230 | 69% | A1B1 | A2B2 | A2B2 | N/A | A2B2 | AF-max + AF-avg | Yes |
| H0217 | 72% | A2B2C2D2E2<br>F2 | N/A | N/A | A2B2C2D2E2<br>F1 | A2B2C2D2E2<br>F1 | TB | No |
| H0258 | 75% | A1B2 | A1B1 | A1B2 | A1B2 | A1B2 | AF-avg + TB | Yes |
| T0218o | 75% | A2 | A3 | A3 | A2 | A2 | TB | Yes |
| H0236 | 76% | A3B6 | A3B3 | A3B3 | N/A | A3B3 | AF-max + AF-avg | Yes |
| T0270o | 78% | A6 | A3 | A3 | A6 | A6 | TB | Yes |
| H0244 | 91% | A2B2C2 | A2B2C1 | A2B2C1 | N/A | A2B2C1 | AF-max + AF-avg | Yes |
| H0267 | 93% | A2B2 | A1B1 | A1B1 | A1B1 | A1B1 | AF-max + AF-avg + TB | Yes |
| H0265 | 100% | A9B18 | A1B1 | A1B1 | N/A | A1B1 | AF-max + AF-avg | No |
| Accuracy on<br>targets applied |  |  | 56%<br>(14/25) | 48%<br>(12/25) | 82.4%<br>(14/17) | 71.4%<br>(20/28) |  | 92.9%<br>(26/28) |
| Accuracy on all 28<br>targets |  |  | 50%<br>(14/28) | 42.9%<br>(12/28) | 50%<br>(14/28) | 71.4%<br>(20/28) |  | 92.9%<br>(26/28) |

On the 28 targets, MULTICOM’s final top-1 prediction was correct for 20 targets, achieving an accuracy of **71.4%**, which highlights its capability to rather accurately predict the stoichiometries of protein multimers. This accuracy significantly outperformed the individual approaches: AF-max (50%), which ranks stoichiometries using AlphaFold3’s maximum ranking score; AF-avg (42.9%), which uses AlphaFold3’s average ranking score; and the template-based (TB) prediction (50%). These results demonstrate that the combination of AlphaFold3-based and template-based predictions in MULTICOM substantially improves accuracy.

It is worth noting that AlphaFold3-based predictions were not applied to three large targets (H0272, H0227, and H0217) because their size exceeded the 5000-residue limit of the AlphaFold3 web server. Additionally, template-based predictions were unavailable for 11 targets, such as H0245 and T0259o, due to the absence of significant complex templates. When evaluated only on targets where each method was applicable, the accuracies of AF-max, AF-avg, and TB increased to 56%, 48%, and 82.4%, respectively. AF-max slightly outperformed AF-avg, although the two methods occasionally succeeded on different targets, such as H0215, T0257o, T0240o, H0229, H0258, and T0259o, showing some complementarity. The much higher precision of template-based predictions underscores their reliability when applicable, highlighting why template-based predictions should take precedence over AlphaFold3-based predictions in cases of inconsistency. However, TB could only be applied to about 61% of the targets (17 out of 28) because some targets do not have significant complex templates, whereas AF-max and AF-avg were applicable to 89% (25 out of 28). This observation highlights the necessity of combining both methods.

The advantage of combining AlphaFold3-based and template-based predictions is evident in three specific situations. **First**, for some of targets without template-based prediction due to lack of significant complex templates, including H0245, T0237o, T0259o, T0240o, T0235o, T0234o, H0229, AF-max and/or AF-avg made correct predictions. **Second**, for two of large targets that AlphaFold3 could not process due to large size (i.e., H0272 and H0227), template-based predictions provided correct results, filling this gap. For another large target H0217, which has a stoichiometry of A2B2C2D2E2F2, a complex template (PDB code: 7F66) was available. This template matched the first five subunits (A2B2C2D2E2) but did not cover subunit F. Theoretically, multiple stoichiometry candidates following the pattern A2B2C2D2E2F? could have been proposed for testing, including A2B2C2D2E2F1 and A2B2C2D2E2F2. However, the correct stoichiometry, A2B2C2D2E2F2 exceeded the 5000 residues limit of AlphaFold3 web server and was not tested. Consequently, the partially correct stoichiometry candidate, A2B2C2D2E2F1, was selected as the final prediction. **Third**, when AlphaFold3-based predictions were incorrect, different template-based predictions rectified the errors for several targets, including H0220, H0233, T0218o, and T0270o, because they were prioritized. With an accuracy of 82.4% on applicable targets, template-based predictions were particularly effective on three hard targets, including H0233, T0218o, T0270o (difficulty value > 60%), where most CASP16 predictors failed.

When the AlphaFold3-based prediction and the template-based prediction differ, in only one case (H0208, stoichiometry: A1B1, a hetero-dimer), the correct AlphaFold3-based prediction (A1B1) was mistakenly replaced by the incorrect template-based prediction (A2B2). The template-based prediction of A2B2 arose because all the complex templates identified for each subunit of H0208 were homodimers, leading to the erroneous conclusion that each subunit must have two copies in the complex. This prediction lacked confidence, as no hetero-multimer template containing both subunits was available. A post-prediction analysis revealed that subunits A and B of H0208 are homologous and can both match the same homodimer template (PDB code: 6UXU). This suggests that a more accurate template-based prediction, hetero-dimer A1B1, can be inferred from this template, considering that homodimers can evolve into hetero-dimers over time. Another challenge associated with this target was the compatibility between A1B1 (hetero-dimer) and A2B2 (hetero-tetramer). The structural models for these two stoichiometries could be superimposed well, creating ambiguity in determining which prediction was correct. This overlap made it particularly difficult to distinguish between the two stoichiometries with confidence. This challenge also caused incorrect predictions for several other hetero-dimer/hetero-tetramer targets.

The AlphaFold3-based methods (AF-max, AF-avg) and the template-based method all produced the same top-1 prediction for six targets (H0225, H0222, H0223, T0206o, H0232, H0267). Among these, five predictions were correct, with only one exception (H0267), resulting in a success rate of 83%. This performance is notably higher than the overall 71.4% accuracy of MULTICOM_AI on all 28 targets. This observation suggests that when AlphaFold3-based predictions (AF-max and AF-avg) and template-based predictions align for a given target, especially one of low or medium difficulty (difficulty value < 50%), the accuracy of the prediction tends to be high. In only one case (i.e., H0267), all the three methods failed in the same manner, predicting the stoichiometry as A1B1 (hetero-dimer) instead of the correct A2B2 (hetero-tetramer). H0267 is the second most difficult target among the 28 targets, with a difficulty value of 93%, meaning that 93% of CASP16 predictors also failed to rank the correct stoichiometry as the top prediction. One reason for this failure is the compatibility between the two stoichiometries, hetero-dimer (A1B1) and hetero-tetramer (A2B2), which made it challenging to confidently identify the correct one. In this case, the prediction incorrectly downgraded the stoichiometry from hetero-tetramer to hetero-dimer, overlooking the possibility that two dimers might oligomerize to form a tetramer.

The accuracy of MULTICOM_AI’s top-3 predictions is **92.9%**, significantly higher than the 71.4% accuracy of its top-1 predictions (**Table 3**). Among the 28 targets, it failed to include the correct stoichiometry within the top three predictions for only two targets, H0217 and H0265. As previously discussed, the failure on H0217 was due to the large size of its correct stoichiometry (A2B2C2D2E2F2), which exceeded the scoring capacity of the AlphaFold3 web server. With the availability of a local installation of AlphaFold3, it is likely that the correct stoichiometry could be included among the top three predictions. For H0265, the correct stoichiometry, a filament with A9B18, fell outside the typical range of subunit copy numbers observed in the templates and was therefore not proposed for AlphaFold3 to rank. This filament target represents the most challenging of all CASP16 Phase 0 targets, as all CASP16 predictors failed to correctly predict its stoichiometry. Predicting the stoichiometry of filaments is particularly difficult because filaments often have a very large and sometimes arbitrary number of subunit copies.

For the remaining 26 targets, the top-3 predicted stoichiometries always included the correct one. This demonstrates that the approach used to propose stoichiometry candidates is highly effective and that AlphaFold3-based predictions can successfully narrow the possibilities to a small set of choices. However, selecting the correct stoichiometry as the top-1 prediction remains a significant challenge.

### 3.3 Analysis of stoichiometry prediction on easy and hard targets

Based on the difficulty metric, the 28 CASP16 targets can be divided into two categories: 14 easy targets with difficulty values less than 60% and 14 hard targets with difficulty values of 60% or higher. For the **14 easy targets** (highlighted with a green background in **Table 2**), MULTICOM_AI achieved a top-1 prediction accuracy of **92.86%** and a top-3 prediction accuracy of **100%**, significantly outperforming its performance on the hard targets, where the respective accuracies were **50%** and **85.7%**. MULTICOM_AI correctly predicted the stoichiometry as the top-1 choice for 13 out of 14 easy targets.

Among these 13 correctly predicted easy targets, the AlphaFold3-based and template-based predictions fully agreed on five targets (H0225, H0222, H0223, T0206o, H0232), with AF-max, AF-avg, and TB all producing the same top-ranked prediction. For two targets (H0215 and T0257o), either AF-max or AF-avg matched the correct prediction made by the template-based method. Additionally, for five targets (H0245, T0259o, T0240o, H0272, and T0237o), one of the methods produced the correct prediction when the other method could not make prediction at all, either due to the absence of templates or the large size of the target. **Figure 3** illustrates how AF-max correctly predicted the stoichiometry (A4) for T0237o in the absence of a template-based prediction. Notably, as the stoichiometry candidate order increased, the ranking score initially rose, peaking at the correct stoichiometry (A4), and then gradually declined.

**Figure 3.**
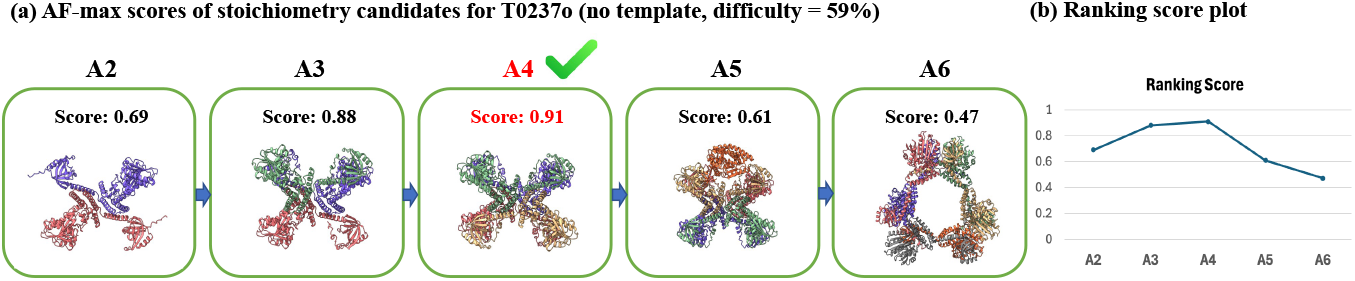
AF-max predicted the correct stoichiometry (A4) for CASP16 target T0237o when there was no template-based prediction. **(a)** AlphaFold3 max ranking scores for stoichiometry candidates (A2-A6); **(b)** The ranking scores are plotted against stoichiometry candidates (from low order to high order).

The template-based prediction (TB) completely conflicted with the AlphaFold3-based prediction (both AF-max and AF-avg) on two easy targets, H0220 and H0208, where the final top-1 prediction based on the template-based method was correct for H0220 (correct stoichiometry: A1B4; **Figure 4**) but incorrect for H0208 (correct stoichiometry: A1B1; **Figure 5**). In the case of H0220, the template information accurately determined the number of copies for subunit B as 4, resulting in the selection of the correct stoichiometry, A1B4, over the A1B1 prediction made by both AF-max and AF-avg.

**Figure 4.**
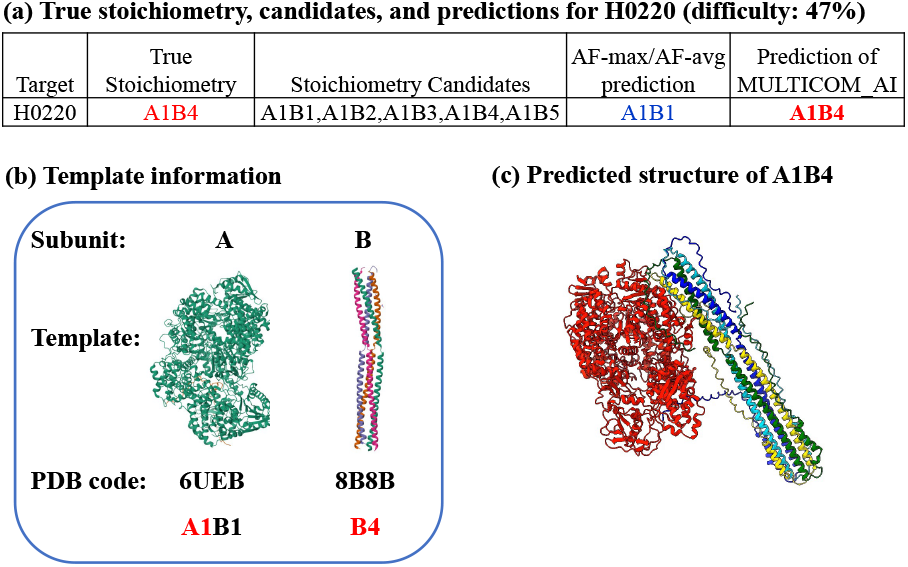
The template-based prediction corrected the error of the AlphaFold3 prediction for H0220. **(a)** The true stoichiometry (A1B4), proposed stoichiometry candidates and top 1 predictions of AF-max / AF-avg and the MULTICOM_AI’s prediction based on the template-based prediction. **(b)** The number of copies of subunits A and B inferred from two templates (PDB codes: 6UEB (stoichiometry of A1B1) and 8B8B (stoichiometry of B4)), respectively. **(c)** The structure of H0220 of the correct stoichiometry A1B4 predicted by AlphaFold3.

**Figure 5.**
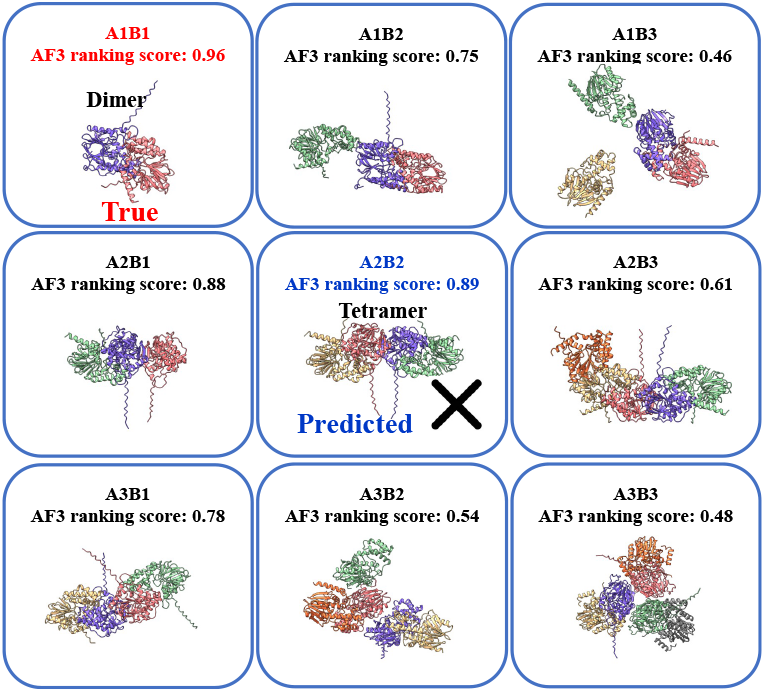
The failure of predicting the stoichiometry (hetero-dimer) of H0208 as hetero-tetramer. According to the AlphaFold3 maximum ranking scores for 9 candidate stoichiometries above, the true stoichiometry (A1B1, hetero-dimer, red) has the highest AlphaFold3 ranking score (0.96), which was correctly selected as top 1 prediction by AF-max. The stoichiometry (A2B2, hetero-tetramer, blue) predicted by the template-based prediction has the second highest score (0.89). The final top 1 prediction of MULTICOM_AI based on the template-based prediction is incorrect.

Conversely, H0208 presented the opposite scenario, where the incorrect template-based prediction (A2B2, hetero-tetramer) overrode the correct AlphaFold3-based prediction (A1B1, hetero-dimer), leading to an overprediction from dimer to tetramer. The compatibility between the hetero-dimer and hetero-tetramer models, combined with their closely ranked maximum scores (0.96 versus 0.89), made it difficult to confidently distinguish between them. As discussed in Section 3.2, the failure of the template-based prediction for H0208 stemmed from the incorrect interpretation of the templates. H0208 was the only easy target for which MULTICOM_AI failed to make correct top-1 prediction.

For the **14 hard targets** (highlighted with an orange background in **Table 2**), MULTICOM_AI achieved correct top-1 predictions for 7 targets, resulting in an accuracy of **50%**. Among these 7 correctly predicted hard targets (H0233, H0258, T0218o, T0270o, T0235o, T0234o, and H0227), the template-based predictions corrected errors in the AlphaFold3-based predictions for four targets (H0233, H0258, T0218o, and T0270o). Meanwhile, the AlphaFold3-based predictions successfully filled gaps where no template-based predictions were available for two targets (T0234o and T0235o). Additionally, for one target (H0227), the template-based prediction provided the correct result when AlphaFold3 could not generate models due to the large size of the target. **Figure 6** demonstrates how the template-based prediction rectified the error in the AF-max prediction for target T0270o (difficulty: 78%).

**Figure 6.**
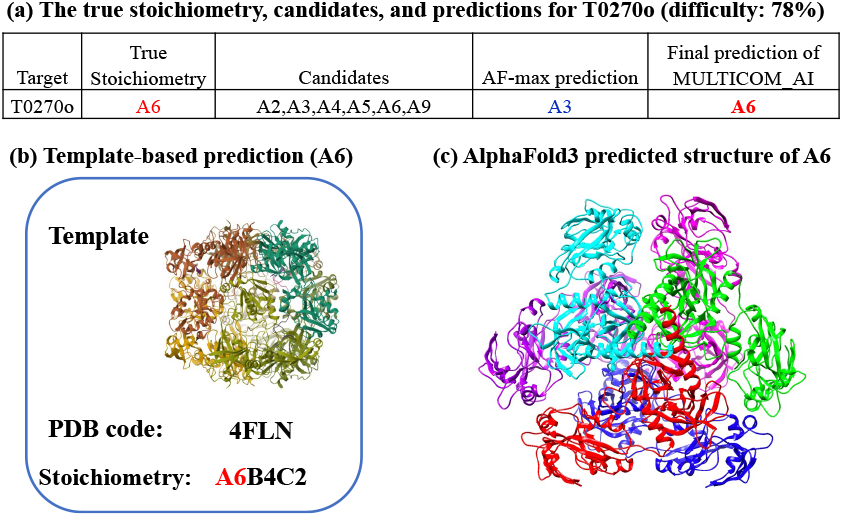
The template-based prediction corrected the error of AlphaFold3 prediction for hard target T0270o (difficulty: 78%). **(a)** The true stoichiometry, proposed stoichiometry candidates, AF-max prediction, and the final prediction of MULTICOM_AI. **(b)** The stoichiometry A6 predicted from a template (PDB code: 4FLN). **(c)** The structure of A6 predicted by AlphaFold3.

MULTICOM_AI failed to make correct top-1 prediction for 7 hard targets (H0229, H0230, H0217, H0236, H0244, H0267, and H0265). Five of them (H0229, H0230, H0236, H0244, and H0265) did not have good templates for template-based prediction, which was one reason they were hard. There are also other factors causing the failure, which are analyzed below.

Both H0229 and H0230 are hetero-dimers, but AF-max and/or AF-avg overpredicted their stoichiometry from hetero-dimer to hetero-tetramer (**Figure 7**), indicating that the compatibility between hetero-dimer and hetero-tetramer makes it hard for AlphaFold3 to distinguish them and is a main source of incorrect prediction. Similarly, as discussed in Section 3.2, the failure on H0267 (a hetero-tetramer) was due to the ambiguity between hetero-tetramer (A2B2) and hetero-dimer (A1B), on which TB, AF-max and AF-avg all underpredicted its stoichiometry as hetero-dimer. It is extremely hard to distinguish tetramer and dimer in this case because all three methods made the same mistake.

**Figure 7.**
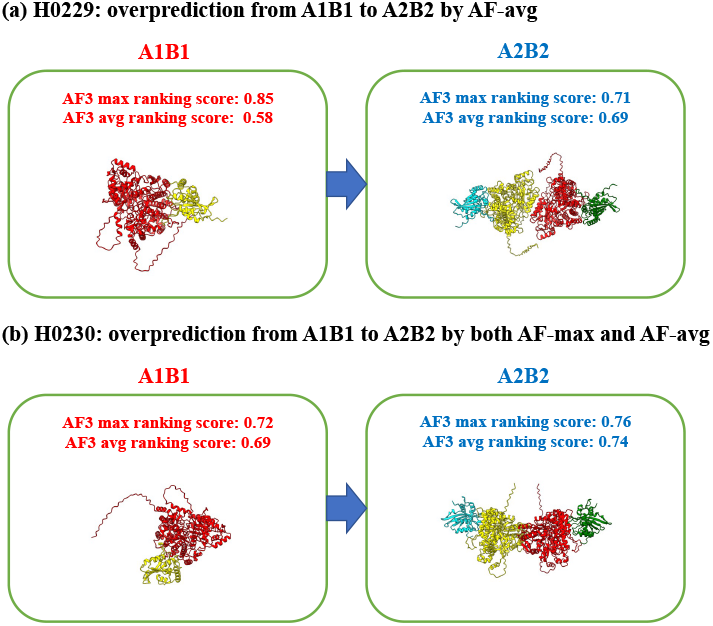
The overprediction of stoichiometry for H0229 and H0230 from hetero-dimer (true) to hetero-tetramer. **(a)** The AF-max and AF-avg scores for stoichiometry A1B1 (hetero-dimer, red) and A2B2 (hetero-tetramer, blue) for H0229. AF-avg incorrectly predicted the stoichiometry as hetero-tetramer, while AF-max correctly predicted hetero-dimer. **(b)** Both AF-max and AF-avg incorrectly predicted the stoichiometry for H0230 as hetero-tetramer. The max or average ranking scores for A1B1 and A2B2 are close, make it difficult to distinguish them.

The failure on H0236 (correct stoichiometry: A3B6) was due to the incorrect stoichiometry ranking of AlphaFold3 and the lack of consideration of the higher-level oligomerization (folding) process. As shown in **Figure 8**, an incorrect stoichiometry (A3B3) has a higher AlphaFold3 maximum ranking score than the correct A3B6, leading to the incorrect prediction (**Figure 8(a)**). However, the stoichiometry of subunits A and B, which were also released as standalone homo-multimer targets T0234o and T0235o separately during CASP16, was correctly predicted as A3 and B6 by AlphaFold3 maximum ranking score respectively (**Figure 8(b)** and **Figure 8(c)**). Considering the hierarchical oligomerization process in which subunits A and B first form A3 and B6 separately and then are assembled into A3B6, the AlphaFold3 prediction error for H0236 could have been corrected. Moreover, the three B subunits in A3B3 models do not interact with each other as the six B subunits in A3B6 models do, where they form a tightly packed 6-unit ring (**Figure 8(a)**), indicating some subunit Bs are missing in A3B3. This example highlights the importance of considering other factors (e.g., higher-order complex oligomerization/folding and the shape and interface of structural models) in addition to the AlphaFold3 ranking scores in selecting stoichiometries for hard targets. H0244 is another case in which the incorrect AlphaFold3-based top-1 prediction may have been corrected by considering another factor – symmetry (**Figure 9**).

**Figure 8.**
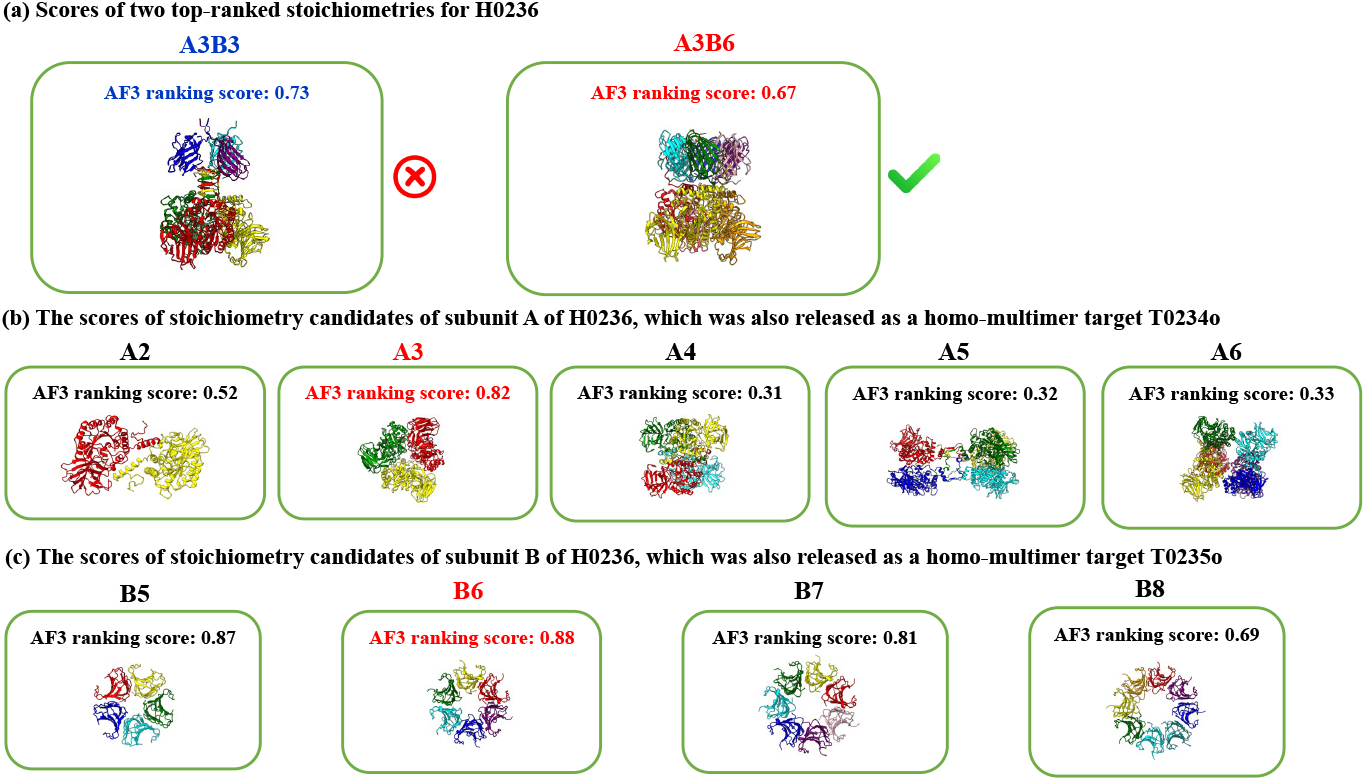
The AlphaFold3 prediction of stoichiometries for three related targets H0236, T0234o, and T0235o. The true stoichiometry of H0236 is A3B6. Its subunits A and B were also released as two separate homo-multimer targets T0234o and T0235o separately. **(a)** The AlphaFold3 maximum ranking scores of the top two ranked stoichiometries (A3B3 and A3B6) for H0236. The incorrect stoichiometry A3B3 has a higher score than the correct one (A3B6), leading to the incorrect prediction. **(b)** The scores of stoichiometry candidates of T0234o (subunit A of H0236). The correct one (A3) has the highest score of 0.82, leading to the correct prediction. **(c)** The scores of stoichiometry candidates of T0235o (subunit B of H0236). The correct one (A6) has the highest score of 0.88, leading to the correct prediction.

**Figure 9.**
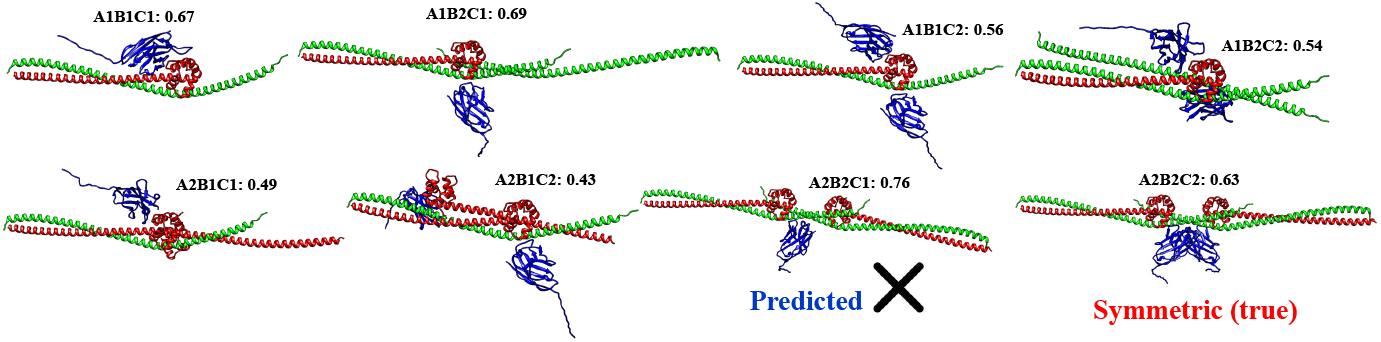
The score of stoichiometry candidates for H0244 (true stoichiometry: A2B2C2; difficulty: 91%). The stoichiometry (A2B2C1) that has the highest AlphaFold3 maximum ranking score is not symmetric, while the stoichiometry (A2B2C2) with the fourth highest score is symmetric. In the asymmetric structural model of A2B2C1, the single copy of subunit C has two identical positions to dock, which is unlikely. If the symmetry were sufficiently considered, A2B2C2 could have been ranked ahead of A2B2C1. However, during CASP16, we only included A2B2C2 into the top three with some consideration of the symmetry.

As for H0265, a filament target that has a very unusual stoichiometry of A9B18, MULTICOM_AI failed to propose the correct stoichiometry as candidate as discussed in Section 3.2, resulting in incorrect top-1 prediction. The failure on H0217 was due to the large size of the true stoichiometry (A2B2C2D2E2F2), which could not be tested by AlphaFold3 webserver and therefore was not evaluated. However, if the symmetry were considered, A2B2C2D2E2F2 may have been preferred over A2B2C2D2E2F1 as top-1 prediction.

H0265 and H0217 are also the only two hard targets that MULTICOM_AI failed to make correct top-3 predictions. MULTICOM_AI made correct top-3 predictions for the other 12 out of 14 hard targets, yielding a high top-3 prediction accuracy of **85.7%**.

### 3.4 Comparison of stoichiometry prediction on homo-multimers and hetero-multimers

In CASP16, MULTICOM_AI achieved a top-1 stoichiometry prediction accuracy of **100% for all 9 homo-multimer targets** (5 easy and 4 hard), compared to **57.9% for 19 hetero-multimer targets** (9 easy and 10 hard) (**Table 2**). These results demonstrate that MULTICOM_AI can predict the stoichiometry of homo-multimers with high accuracy, whereas predicting the stoichiometry of hetero-multimers is significantly more challenging.

One primary reason for this disparity is that homo-multimers inherently have fewer potential stoichiometry candidates (e.g., A2 to A9) compared to the vast combinations possible for hetero-multimers, which involve multiple unique subunits and varying copy numbers. The smaller candidate pool for homo-multimers increases the likelihood of including the correct stoichiometry in the candidate list and ranking it at the top. Indeed, for all 9 homo-multimer targets, the true stoichiometry was included in the candidate list. Among these, AF-max selected the true stoichiometry as the top prediction for 6 targets, achieving an accuracy of 66.67%, which is substantially higher than its 50% accuracy for applicable hetero-multimers. Similarly, AF-avg correctly identified the true stoichiometry for 5 homo-multimer targets, yielding an accuracy of 55.56%, compared to 43.8% for applicable hetero-multimers. When considering the best predictions of both AF-max and AF-avg collectively, the true stoichiometry was ranked as the top prediction for 7 out of 9 homo-multimer targets, resulting in an overall accuracy of 77.8%.We hypothesize that AlphaFold3’s superior performance on homo-multimers is also due to their greater structural symmetry and lower complexity compared to hetero-multimers, which often feature more intricate interactions among diverse subunits.

For the two homo-multimer targets (T0218o and T0270o) where both AF-max and AF-avg failed to rank the true stoichiometry as the top prediction, the template-based method successfully rescued the prediction. In these cases, significant complex templates were available. For example, as illustrated in **Figure 10**, a template (PDB code: 4W8J) with stoichiometry A2 was used to correctly predict the stoichiometry of T0218o when AF-max and AF-avg incorrectly selected A3.

**Figure 10.**
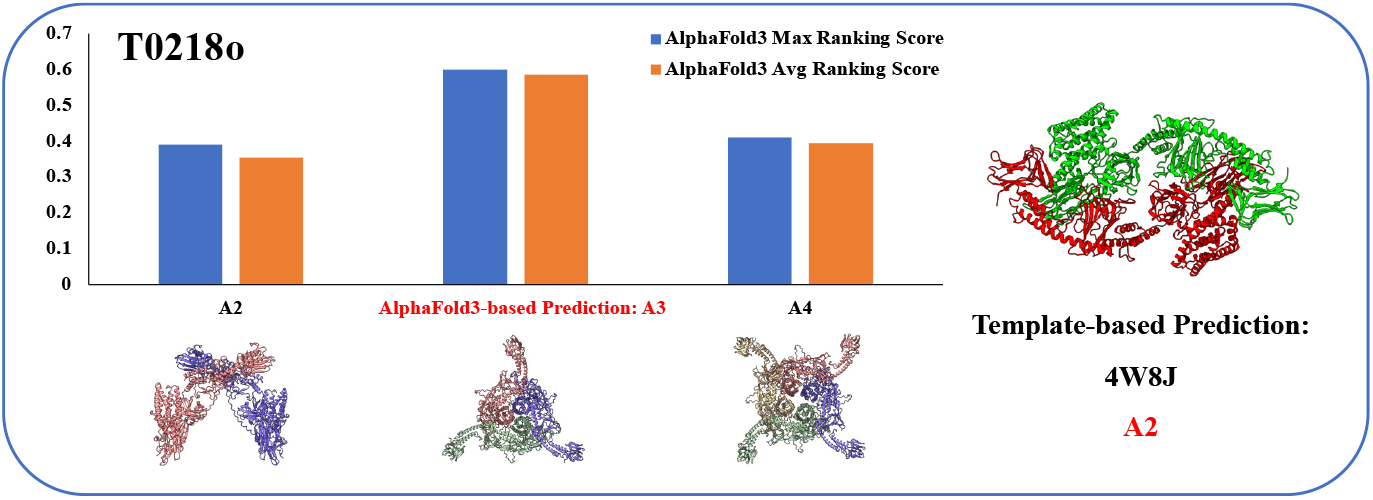
The AlphaFold3 prediction versus the template-based prediction for T0218o. AF-max and AF-avg incorrectly ranked stoichiometry A3 at the top, but the template-based approach correctly predicted stoichiometry A2 using the stoichiometry of a template (PDB code: 4W8J, A2) with its two chains coloured in green and red.

Additionally, homo-multimers consist of only one unique subunit, which increases the likelihood of finding appropriate complex templates compared to hetero-multimers, which require templates that encompass multiple unique subunits. This further explains why predicting the stoichiometry of homo-multimers is inherently easier than that of hetero-multimers for our approach. Different from the perfect prediction accuracy of our approach on the homo-multimer targets, many CASP16 predictors failed on some of them, indicated by the high difficulty value of some homo-multimer targets (e.g., T0270o, difficulty = 78%).

The **accuracy of the top-3 predictions for 19 hetero-multimers is 89.5%**, significantly higher than the **57.9% accuracy of top-1 predictions**. The top-3 predictions failed to include the true stoichiometry for only two targets, H0217 and H0265, because their correct stoichiometry was not proposed at all. For H0217, this was due to its large size, and for H0265, it was due to its large and unusual stoichiometry (A9B18), as discussed in Section 3.2.

For the remaining 17 hetero-multimer targets, the true stoichiometry was successfully included in the top 3 predictions. However, MULTICOM_AI was unable to rank the true stoichiometry as the top choice for six of these targets (H0208, H0229, H0230, H0236, H0244, H0267), underscoring the challenge of accurately identifying the correct stoichiometry among the top three candidates. As outlined in Sections 3.2 and 3.3, one major difficulty stems from the ambiguity and compatibility between hetero-dimers and hetero-tetramers, which can overlap structurally. This structural similarity between their models often makes it difficult for the AlphaFold3 ranking score to differentiate and rank them accurately.

Four of the failed cases (H0208, H0229, H0230, and H0267) illustrate this challenge. For H0208, H0229, and H0230, as shown in **Figures 5** and **7**, the structural models for the two stoichiometries (A1B1 and A2B2) are fully superimposable (**Figure 11**). In the case of H0267, all three methods (AF-max, AF-avg, and TB) were similarly confused by the structural ambiguity between dimers and tetramers, resulting in the same incorrect prediction. It is worth noting that some protein targets, such as these, might naturally exist in multiple stoichiometries with different proportions in cells. There is a possibility that experimental structure determination captures only one stoichiometry, while computational predictions may reveal multiple possibilities.

**Figure 11.**
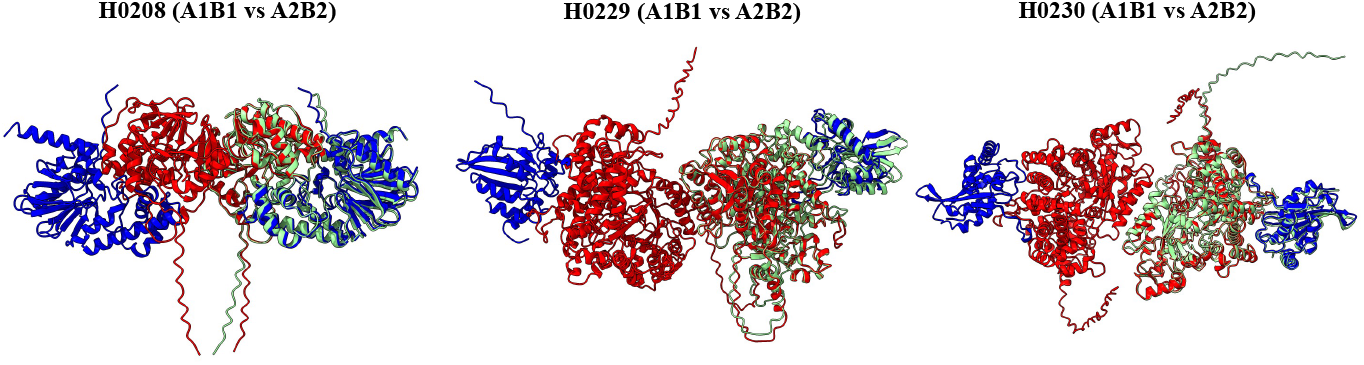
Superimposition of structural models for two stoichiometry candidates, A1B1 and A2B2, for three hetero-dimer targets (H0208, H0229, H0230) with a true stoichiometry of A1B1. In the A2B2 models, the two subunits are represented in blue and red, while the A1B1 models are shown in green. The A1B1 models align with one-half of the A2B2 models, highlighting their partial overlap and the difficulty of differentiate them.

The final two targets among the six where MULTICOM_AI failed in top-1 predictions are H0236 and H0244, both of which lacked significant templates. However, as illustrated in **Figures 8** and **9**, if additional considerations, such as higher-order oligomerization and symmetry, had been factored into the predictions, the true stoichiometry for these targets could have been ranked at the top.

Among the 16 hetero-multimer targets with sequence lengths below 5000 residues AlphaFold3 was applied to, AF-max correctly selected the true stoichiometry as the top-1 prediction for 8 targets, achieving an accuracy of 50%. In comparison, AF-avg correctly identified the true stoichiometry for 7 targets, yielding an accuracy of 43.8%.

For instance, in the case of target H0245, both AF-max and AF-avg assigned a higher ranking score to the correct stoichiometry (A1B1) than to other alternative candidates (A1B2, A2B1, and A2B2) proposed from template information (**Figure 12**). Notably, H0245 lacked a complex template encompassing both subunits, and therefore, no template-based prediction was available for this target. This highlights the capability of AlphaFold3 to accurately predict stoichiometry even in the absence of comprehensive template information.

**Figure 12.**
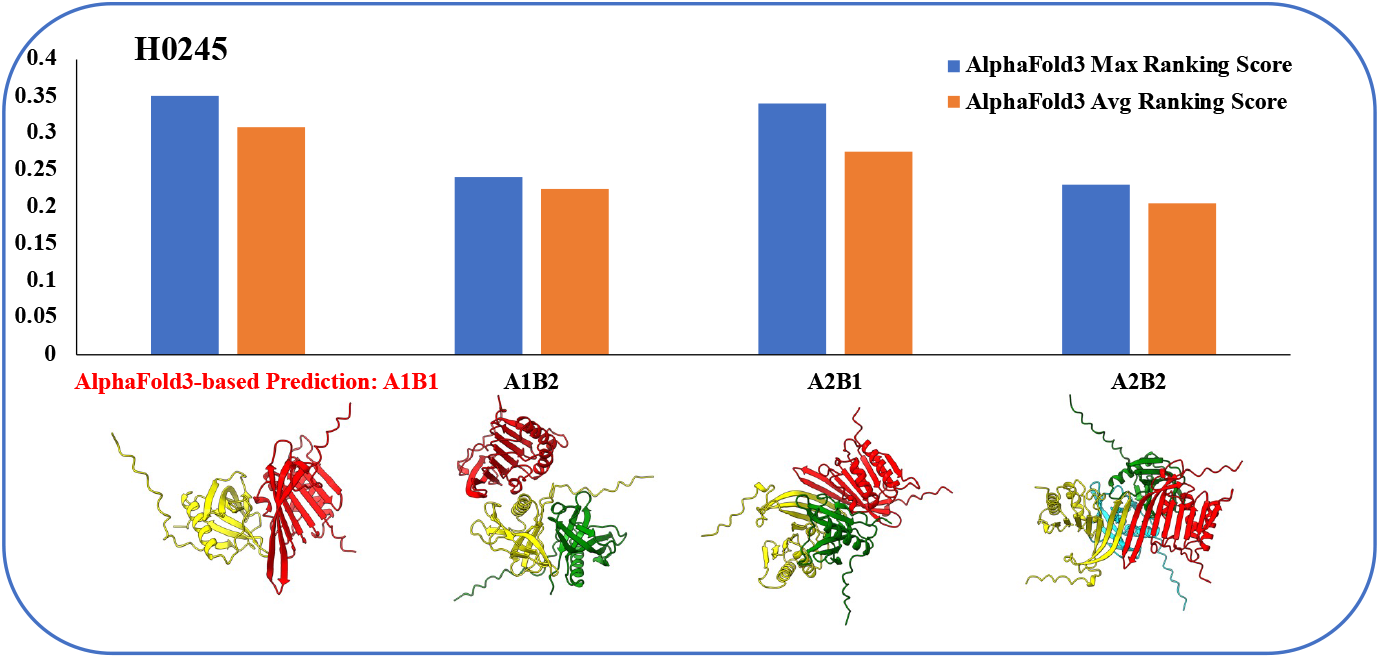
The AlphaFold3 ranking scores for four stoichiometry candidates for H0245. The true stoichiometry (A1B1) has the highest maximum or average score.

However, the overall top-1 prediction accuracy for hetero-multimers remains at just **50%** for AF-max, underscoring the need for significant improvements in distinguishing the correct stoichiometry among competing candidates.

In contrast, template-based predictions were made for 13 hetero-multimers, yielding 10 correct top-1 predictions and achieving an accuracy of **76.9%**. Among the 10 targets with both AlphaFold3-based and template-based predictions, the template-based approach successfully corrected errors in AlphaFold3 predictions for two targets: H0220 (**Figure 4**) and H0233.

For H0233, it was observed that two copies of a homologous template for subunit A were present in complex template 7RK1, two copies of a homologous template for subunit B were present in complex template 8SZY, and two copies of a homologous template for subunit C were present in complex template 7CAI. This evidence suggested that each subunit of H0233 exists in pairs, leading to the accurate prediction of its stoichiometry as A2B2C2 (**Figure 13**). The template-based prediction corrected the erroneous stoichiometry ranking from AlphaFold3 for this target, underscoring the value of integrating template-based and AlphaFold3-based approaches for stoichiometry prediction.

**Figure 13.**
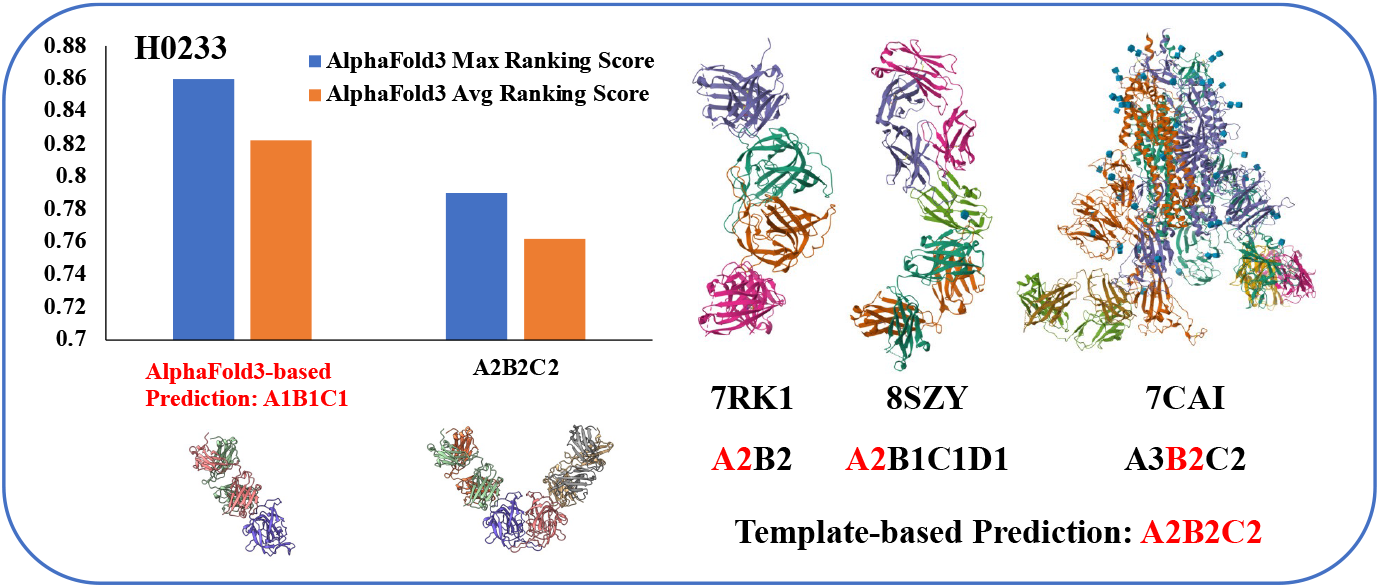
Template-based prediction versus AlphaFold3-based prediction for H0233. The template-based approach found three templates (7RK1, 8SZY, and 7CAI) for three subunits (A, B, C) for H0233 respectively, each of which contains two copies of the templates for each subunit, leading to the correct prediction of stoichiometry A2B2C2 for H0233. However, the AlphaFold3-based methods ranked the incorrect stoichiometry A1B1C1 ahead of A2B2C2. MULTICOM_AI chose the template-based prediction as top 1.

## 4. Discussion

Our experiment demonstrates that using template information or simple enumeration to generate a shortlist of stoichiometry candidates is highly effective, as the proposed stoichiometries include the true ones for 26 out of 28 targets (**sensitivity: 93%**). The two exceptions, H0217 and H0265, were missed due to limitations imposed by AlphaFold3 web server’s protein length restriction (5000 residues) and the unusually large stoichiometry of H0265 (A9B18). The release of AlphaFold3 software is expected to address the length limitation, as locally installed versions with substantial GPU memory should be capable of handling larger proteins. However, proposing and testing very large stoichiometries, such as A9B18, will remain challenging without template information due to the vast number of candidates, which makes accurate selection difficult.

Given a shortlist of stoichiometry candidates, AlphaFold3 ranking scores, particularly the maximum ranking scores (AF-max), are highly effective in identifying the top three stoichiometries. For 24 CASP16 targets where the true stoichiometry was proposed and AF-max was applied, AF-max ranked the true stoichiometry among the top three for 23 targets (**accuracy: 95.8%**), missing only H0244. However, ranking the true stoichiometry as the top-1 prediction remains challenging. AF-max identified the true stoichiometry as the top-1 choice for 14 out of 24 targets (**accuracy: 58%**), which is 38 percentage points lower than its top-3 accuracy, leaving significant room for improvement.

The most effective way to correct errors in the AlphaFold3-based stoichiometry ranking is through template-based predictions inferred from significant complex templates when available. Template-based predictions are more accurate at selecting the top-1 stoichiometry than AlphaFold3 ranking scores. Among 14 targets with both AlphaFold3-based and template-based predictions, the template-based predictions achieved an accuracy of **85.7%** (12 out of 14 targets), compared to **64.29%** (9 out of 14 targets) for AlphaFold3-based predictions. Consequently, MULTICOM successfully used template-based predictions to correct AlphaFold3 ranking errors for all but one target they disagreed on (e.g., H0220, H0233, T0218o, and T0270o). The exception, H0208 (a hetero-dimer), failed due to incorrect template interpretation and the ambiguity between hetero-dimer and hetero-tetramer. Template-based predictions are also crucial for very large targets, such as H0272, where AlphaFold3-based predictions cannot be applied.

Despite its utility, template-based predictions could only be confidently applied to 61% of CASP16 targets (17 out of 28), limiting their scope. For the remaining 39% of targets (11), which lacked significant complex templates, AlphaFold3 predictions were indispensable. Among these 11 targets, the AlphaFold3-based approach achieved an accuracy of 54.5% for top-1 predictions, correctly identifying the true stoichiometry for 6 targets. This performance is comparable to its overall accuracy of 58.3% across all 24 targets where true stoichiometries were proposed, indicating that AlphaFold3’s effectiveness is largely independent of template availability and thus robust.

The strong complementarity between AlphaFold3-based and template-based predictions allowed MULTICOM to achieve a good top-1 prediction accuracy of **71.4%** and an impressive top-3 accuracy of **92.9%** across all 28 targets. **Figure 14** illustrates how the two AlphaFold3-based prediction methods (AF-max and AF-avg) and the template-based method complement each other, resulting in improved final predictions.

**Figure 14.**
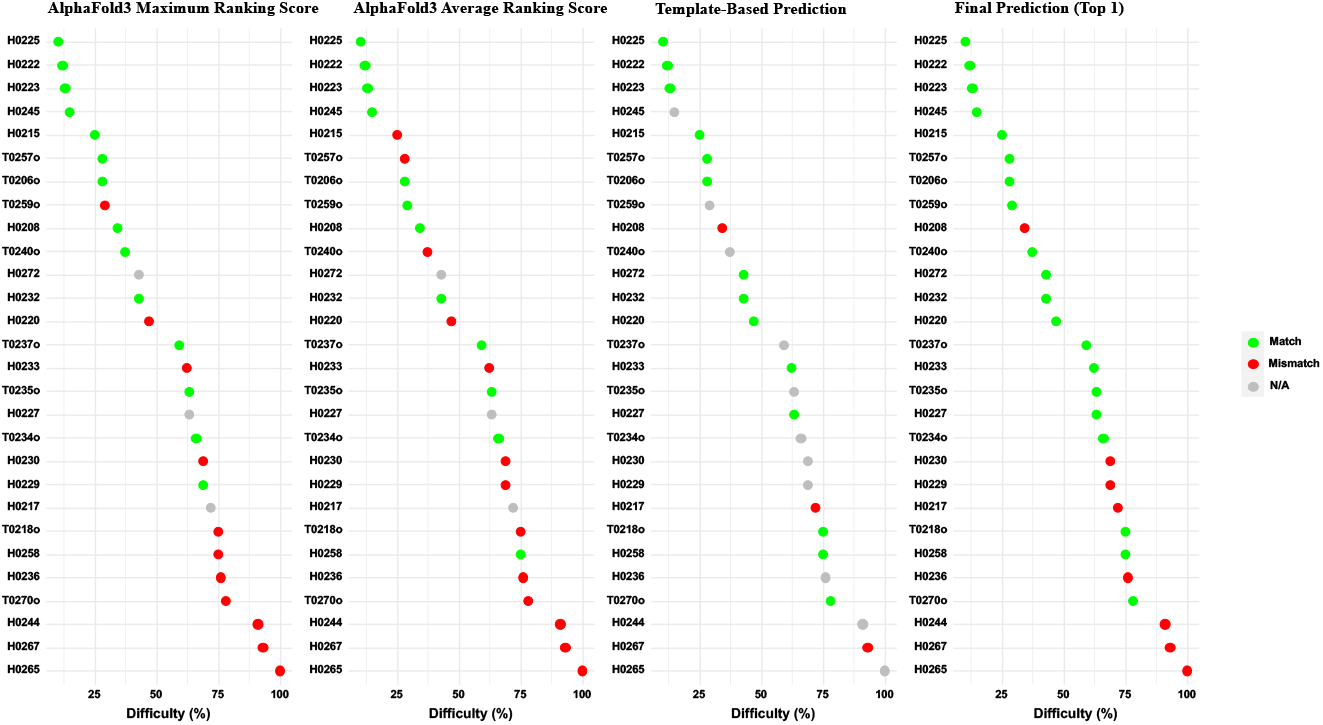
The complementarity of the AlphaFold3-based prediction (AF-max and AF-avg) and the template-based prediction on 28 CASP18 multimer targets. The targets are ordered from low difficulty to high difficulty. Green dots denote correct top-1 predictions, grey dots missing predictions, and red dots incorrect top-1 prediction. The final prediction benefits from the AlphaFold3-based prediction and the template-based prediction correctly fill in some of each other’s missing predictions. The template-based prediction also helps correct several errors made by the AlphaFold3-based prediction when their predictions are different.

A key challenge for AlphaFold3-based predictions in selecting the true stoichiometry as the top-1 choice lies in the ambiguity or similarity between compatible stoichiometries, particularly between hetero-dimer and hetero-tetramer configurations. This issue is evident in three of the five targets without template-based predictions where AlphaFold3 failed (i.e., H0229, H0230, and H0267). To improve prediction accuracy for such cases, additional information needs to be identified and incorporated into the prediction process in the future.

For the other two targets without significant complex templates (i.e., H0236 and H0244), where AlphaFold3-based predictions also failed to make correct top-1 predictions, we observed that other factors such as structural symmetry, structural interfaces, and higher-order oligomerization could potentially correct these errors (**Figures 8 and 9**). However, systematically and automatically leveraging this information remains a significant challenge.

While our combined approach has achieved promising accuracy in stoichiometry prediction, the template-based prediction process still involves manual steps. These include inferring stoichiometry from partial complex templates that do not fully cover all the subunits of a target and analyzing related literature. Automating this process will require advanced machine learning and AI methods capable of integrating diverse sources of information. Developing such tools to streamline and enhance the template-based prediction process represents a crucial next step for improving the overall accuracy and efficiency of stoichiometry predictions.

In evaluating stoichiometry predictions, we defined accuracy based on exact matches with the true stoichiometry, ignoring similarities between stoichiometries. However, some stoichiometries (e.g., A1B1 and A2B2 for H0208 and H0229) produce structurally similar models, while others (e.g., A3 and A6 for T0234o) differ significantly. Thus, exact matches alone may not fully reflect prediction quality.

An alternative is using structural metrics like QS-score^30^ and TM-score TM-score^31^ of structural models to assess the similarity between predicted and true stoichiometries. These metrics are especially relevant when the predicted stoichiometry is a subset (e.g., H0267) or superset (e.g., H0208) of the true stoichiometry, common in higher-order oligomerization. For example, if A1B1 is predicted instead of A2B2 or vice versa, QS-score and TM-score can still quantify partial correctness. In CASP16, such cases were common, particularly for hetero-dimers and hetero-tetramers (e.g., H0208, H0229, H0230, H0267). For another case H0244, MULTICOM_AI predicted A2B2C1 instead of A2B2C2 (true stoichiometry), yet the TM-score of 0.625 of a structural model built for A2B2C1 indicated significant similarity between A2B2C1 and the true stoichiometry.

While QS-score and TM-score are valuable for evaluating compatibility and similarity, they depend on structural model quality, which isn’t always better for closer stoichiometries. Therefore, these metrics should complement, not replace, exact match evaluations.

## 5. Conclusion

In this work, we developed an innovative and effective approach that combines protein template information with AlphaFold3-based complex structure prediction to accurately predict the stoichiometry of protein complexes. The method was blindly tested in the 2024 community-wide CASP16 experiment, where it demonstrated outstanding performance. It consistently generated a shortlist of candidate stoichiometries containing the correct one for almost all tested targets and ranked the correct stoichiometry as the top-1 choice with good accuracy and within the top-3 with high accuracy. This approach addresses a significant gap in the availability of accurate methods for predicting protein stoichiometry—an important yet understudied problem. It provides a powerful tool for scientists to investigate the stoichiometry of many uncharacterized protein complexes lacking prior stoichiometry information, thereby enabling the application of complex structure prediction to them.

Future work will focus on improving the approach further by identifying and leveraging additional sources of information, such as symmetry, structural and interface features, and literature data, through machine learning and artificial intelligence. These enhancements aim to reduce errors in selecting top-1 predicted stoichiometry, as well as to fully automate and scale up the prediction process.

## Acknowledgement

This work was supported by two NIH NIGMS grants (R01GM093123 and R01GM146340) and one NSF grant (DBI2308699).

## Data Availability

The compiled CASP16 stoichiometry prediction dataset is available at: https://github.com/jianlin-cheng/prestoi/tree/main/CASP16_Phase_0_dataset.

## Code Availability

The source code of AlphaFold3-based stoichiometry ranking is available at: https://github.com/jianlin-cheng/prestoi.

