## Supplementary material for "Accurate Prediction of Protein Complex Stoichiometry by Integrating AlphaFold3 and Template Information": Supplmentary material

**Table S1.** The stoichiometry prediction accuracy of the 21 CASP16 predictors on 28 Phase 0 targets. Bold denotes the highest accuracy. MULTICOM_AI ranked at top in terms of the accuracy of both top-1 prediction and all-predictions.

| Predictor | Accuracy of top-1 prediction | Accuracy of all-predictions | Average |
|---|---|---|---|
| MULTICOM_AI | **0.714** | **0.929** | **0.8215** |
| NKRNA-s[1] | **0.714** | 0.786 | 0.75 |
| Zheng-Multimer[2] | 0.679 | 0.714 | 0.6965 |
| Schneidman[3] | 0.679 | 0.714 | 0.6965 |
| Zheng[4] | 0.679 | 0.679 | 0.679 |
| MIEnsembles-Server[5] | 0.643 | 0.750 | 0.6965 |
| CSSB_experimental[6] | 0.643 | 0.714 | 0.6785 |
| CSSB-Human[7] | 0.643 | 0.714 | 0.6785 |
| GromihaLab[8] | 0.643 | 0.643 | 0.643 |
| KiharaLab[9] | 0.607 | 0.750 | 0.6785 |
| OpenComplex[10] | 0.500 | **0.929** | 0.7145 |
| OpenComplex_Server[10] | 0.500 | 0.893 | 0.6965 |
| MultiFOLD2[11] | 0.500 | 0.500 | 0.5 |
| McGuffin[12] | 0.500 | 0.500 | 0.5 |
| elofsson[13] | 0.429 | 0.643 | 0.536 |
| PEZYFoldings[14] | 0.429 | 0.464 | 0.4465 |
| AF3-server[13] | 0.393 | 0.536 | 0.4645 |
| kiharalab_server[9] | 0.286 | 0.286 | 0.286 |
| APOLLO[15] | 0.250 | 0.250 | 0.25 |
| ARC[16] | 0.143 | 0.250 | 0.1965 |
| COAST[15] | 0.107 | 0.179 | 0.143 |

**Table S2.** The stoichiometry predictions of MULTICOM_AI for the 28 Phase 0 targets. Bold denotes the true stoichiometry.

| Target Name | Top 1 Prediction | Top 2 Prediction | Top 3 Prediction | Top 4 Prediction | Top 5 Prediction |
|---|---|---|---|---|---|
| H0208 | A2B2 | **A1B1** | A2B1 | A3B1 | A3B3 |
| H0215 | **A1B1** | A2B2 | A2B1 | | |
| H0217 | A2B2C2D2E2F1 | | | | |
| H0220 | **A1B4** | A1B1 | | | |
| H0222 | **A1B1C1** | | | | |
| H0223 | **A1B1C1** | | | | |
| H0225 | **A1B1C1** | | | | |
| H0227 | **A1B6** | | | | |
| H0229 | A2B2 | **A1B1** | | | |
| H0230 | A2B2 | **A1B1** | | | |
| H0232 | **A2B2** | | | | |

| | | | | |
|---|---|---|---|---|
| H0233 | **A2B2C2** | A1B1C1 | | |
| H0236 | A3B3 | **A3B6** | | |
| H0244 | A2B2C1 | A1B1C1 | **A2B2C2** | A1B2C1 |
| H0245 | **A1B1** | | | |
| H0258 | **A1B2** | A1B1 | | |
| H0265 | A1B1 | **A2B2** | | |
| H0267 | A1B1 | A4B4 | **A2B2** | |
| H0272 | **A1B1C1D1E1F1G1H1I1** | | | |
| T0206o | **A2** | A4 | A6 | A8 |
| T0218o | **A2** | A3 | | |
| T0234o | **A3** | | | |
| T0235o | **A6** | A5 | | |
| T0237o | **A4** | | | |
| T0240o | **A3** | | | |
| T0257o | **A3** | | | |
| T0259o | **A3** | | | |
| T0270o | **A6** | A3 | | |
